# Estimation of motion direction and speed using an organic-semiconductor retinal prosthetic in a blind retinae

**DOI:** 10.64898/2026.04.23.720306

**Authors:** Abhijith Krishnan, C S Deepak, K S Narayan

## Abstract

For a vision system, estimating the speed and direction of movement at the retinal input stage is an essential function for survival in many organisms. Retinal ganglion cells specific to this movement function were identified using multi-electrode array recordings in neonatal chick retina. Motion-evoked “visual streaks” and direction selective responses were observed in chick ganglion cells upon sequential activation as a response to moving bar stimuli. These characteristics were preserved in the sub-retinal prosthetic consisting of a semiconductor polymer film coupled to the blind chick retina which generated spatiotemporal activity patterns resembling those in natural vision. The motion parameters of direction and speed inferred from these recordings demonstrate that polymer-based prostheses can evoke physiologically relevant activity patterns, suggesting their potential to restore motion perception in degenerative retinae.

## 1 Introduction

Estimation of the speed and direction of moving objects is a crucial feature in vision and has a direct role in a natural environment, such as in a predator-prey scenario. The visual signals from the retina need to be processed rapidly to quickly ascertain and identify the speed and direction of moving object in a static or changing background[1]. These survival mechanisms have contributed to the evolution of the retina and the functionality of RGCs[2]. It has been noted that across many species, especially in the avian family that a direction-sensitive mechanism is present upfront in the sensory system. Besides identifying these type of specific retinal neurons in a normal postnatal chick retina, it will also be significant to see these features preserved when artificial photoreceptors are introduced in a blind retina. Thus, it is of utmost importance that any prosthetic device can generate activity patterns in the retina that can reliably transmit information regarding object motion for further processing to the brain.

There are two main ways by which the retina estimates the direction of moving objects. The first uses a specialized subset of retinal ganglion cells (RGCs); Direction Selective RGCs (DSRGC), that exhibit strong (spiking) response to motion in one particular (preferred) direction as opposed to a weak response in the opposite (null) direction[3]. This breaking of translational symmetry in the detection scheme with respect to motion by a single neuron (DSRGC) arises from the upstream input network from starburst amacrine cells[4]. This is thought to be the main pathway for direction estimation in non-primate mammals and other vertebrates, such as the chicken. Recent studies have also shown the presence of such DSRGCs in primates, although the percentage of such cells is lower in primates compared to other mammals[5]. The other mechanism is the use of a “visual streak” caused by the movement of an object on the retina, similar to that in a streak camera. As the object moves over the receptive field of an RGC, it induces firing, which leaves a changing firing pattern (visual streak) on the retina that the brain can use to infer the speed and direction of the moving object. This activity pattern is from a population of different cell types and thus may lead to a less noisy estimation of direction[6]. These methods of motion detection work in tandem with head/eye movements to determine the direction and speed of the object[7]. Although this type of work on DSRGCs[8] and the visual streak[9, 10] for direction estimation has been done in many animals, reports of such features and mechanisms in the chick retina are not available. The chick retina provides a model system for exploring retinal network development and for examining key features of vision, such as colour processing and the segregation of visual information into ON and OFF channels[11]. Identification of direction-selective retinal ganglion cells (DSRGCs) in the chick retina would therefore be informative for understanding the development and functional organization of vision in avian species[12].

The developing chick retina, prior to the formation of photoreceptor outer segments and synapses with bipolar cells, has been used in our laboratory as a blind system that serves as an effective model for a degenerative retina[13, 14, 15]. Previous work on reproducing the visual streak using a retinal prosthesis has been done using multi-electrode arrays (MEA) interfaced epiretinally with the retina to deliver current injections at individual RGC resolutions[16]. This solution, while able to reproduce the firing patterns of RGCs, requires the introduction of an externally driven, highly dense prosthetic device that needs to be surgically inserted epiretinally. In this study, we demonstrate that a subretinally placed, coarsely patterned photoactive polymer layer can elicit visual streaks similar to those observed in the neonatal retina.

In an earlier work, we demonstrated that the temporal response characteristics of a bulk-heterojunction polymer (BHJ) blend layer coupled to a blind retina exhibit features similar to those observed in the neonatal retina[15]. In the present study, we expand on these results by probing the encoding of stimulus direction and speed. Retinae from neonatal and developing chicks were interfaced with MEAs to record action potentials from RGCs. Control studies were performed on neonatal chick retinae to determine how the direction and speed of the object is encoded in a healthy retina. Cells were classified as ON, OFF or ON-OFF based on their response to flashes of light[17]. Bars of light moving in different directions and speeds (+x, -x, +y and -y) were used as stimuli to estimate motion direction and speed. Direction and speed were inferred from the spatiotemporal ‘visual streak’ formed by peak firing times across electrodes. The same stimulus also revealed the presence of direction-selective RGCs (DSRGCs). Patterned BHJ coated prosthetics were coupled subretinally with a developing chick embryo and the same visual stimuli were provided. In the polymer-coupled embryonic retinae, we observed similar visual streaks, from which motion direction and speed could likewise be recovered.

## 2 Results

Multi-electrode recordings from RGCs were obtained from chick retinae at different developmental stages using a 60-electrode MEA with a 200 µm inter-electrode distance, connected to a MEA-2100 amplifier [18]. Control recordings were taken from postnatal (P0–P4) chicks. Neonatal chick retinae were placed with the RGC-side interfacing the MEA to record action potentials. Light stimuli were presented from the bottom surface of the MEA, reproducing the physiological condition in which incident light passes through all retinal layers before reaching the photoreceptors (Figure 1 a). Responses of individual RGCs were extracted from the multi-unit activity through spike sorting [19].

**Figure 1:**
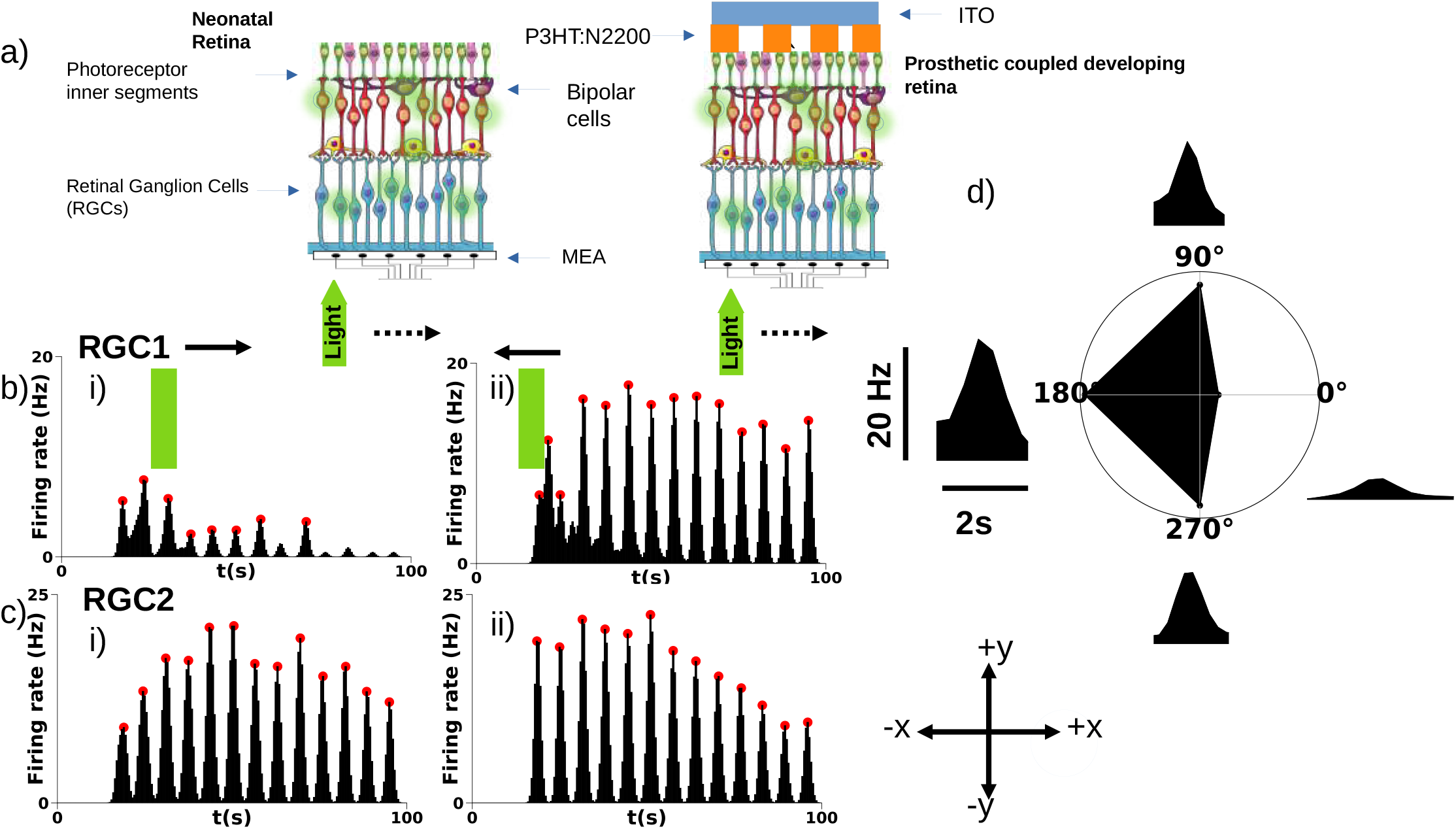
Response of a direction selective RGC (RGC1) and a non-direction selective RGC (RGC2) to object motion in +x and -x directions. a) Schematic of the recording setup with the retina placed RGC side down on the MEA. Light stimulus was provided from the bottom. b) Example of direction selective responses from RGC1 to i)+x and ii)-x direction object motion. c) Example of non-direction selective responses from RGC2 to i)+x and ii)-x direction object motion. Each peak in the firing rate (red points) corresponds to the response of the RGC to a single trial of the bar stimulus. d) Polar plot showing the trial averaged response of RGC1 to object motion in the four directions.

### 2.1 Estimation of direction and speed from the neonatal retina

Full-field flashes of light were used to assess the RGC response features. Response polarity was determined from firing rates evoked during a 500 ms flash of light, using the ratio of peak firing rates during light ON (*R*_*ON*_) versus light OFF (*R*_*OFF*_) periods[17] (Supplementary Fig. 1). Bars of light (200 µm width) moving uniformly across the MEA in different directions were used as stimuli to characterize RGC responses to object motion. Evidence of direction-selective RGCs in the chick retina was obtained by studying and analysing the response of individual RGCs to the moving bar stimulus. The average response to this stimulus carried out over twenty-five cycles, clearly uncovered DSRGCs that repeatedly exhibited asymmetric firing responses to bar movement. The responses of two RGCs to motion in the +x and -x directions are shown in Figure 1. Object motion in the null direction (+x) (Figure 1 b)i)) elicited weak responses from RGC1 and motion in the preferred direction (-x) elicited strong responses (Figure 1 b)i)). Each peak represents the response to a single trial of bar movement over the RGC. Non direction selective RGCs showed similar responses to stimuli moving in both directions (Figure 1 c)i) and ii)). The polar plot (Figure 1 d)) shows the average response of RGC1 to bar motion in the four directions. These cells typically showed firing during both ON and OFF periods to the full-field flash stimulus (Supplementary Figure 2). The collective responses from a population of four types of these DSRGCs tuned to motion in the four cardinal directions can be used by downstream neurons to arrive at an accurate perception of the moving object.

#### 2.1.1 Visual streak from a population of RGCs

In addition to the DSRGCs mentioned above, RGCs in the chick retina showed a visual streak of activity due to the moving bar stimulus. When the bar passed over the receptive field of an RGC [20], the cell fired a burst of spikes, thus generating a streak of activity across the recorded region as the bar traversed successive RGC receptive fields. This spatiotemporal pattern can be used to construct an activity map to estimate the direction and speed of the moving object. Cells showing direction selective responses were excluded from the visual streak based motion calculations since their responses differ for motion in opposite directions. There are multiple ways by which the brain could use RGC responses to infer the direction and speed of a moving object. One such approach relies on the timing of the first spike from individual RGCs[21, 22]. This assumes that RGCs fire with millisecond-scale temporal precision. However, in our recordings, chick RGCs exhibited spike-timing jitter on the order of tens to hundreds of milliseconds, making the time of first spike unreliable for motion estimation. Previous work on motion estimation in primates suggests the need for more than one spike for accurate estimation of the direction and speed of the object[9, 10]. Hence, in this study, we binned spikes from individual RGCs to calculate firing rates and used the time of peak firing to estimate the direction and speed of the stimulus. This RGC response binning procedure is equivalent to a low-pass filtering process, which is typically performed by downstream visual processing regions of the brain.[23]. By plotting the raster for multiple trials of the moving bar stimulus, temporal differences in firing onset for RGCs at different spatial locations were observed(Figure 2 a)). Due to the lower electrode density of the MEA, it was not possible to extract electrical images for individual RGCs[24]. Hence, the spatial location of each RGC was defined as the position of the electrode that recorded its response (Figure 2 b)).

**Figure 2:**
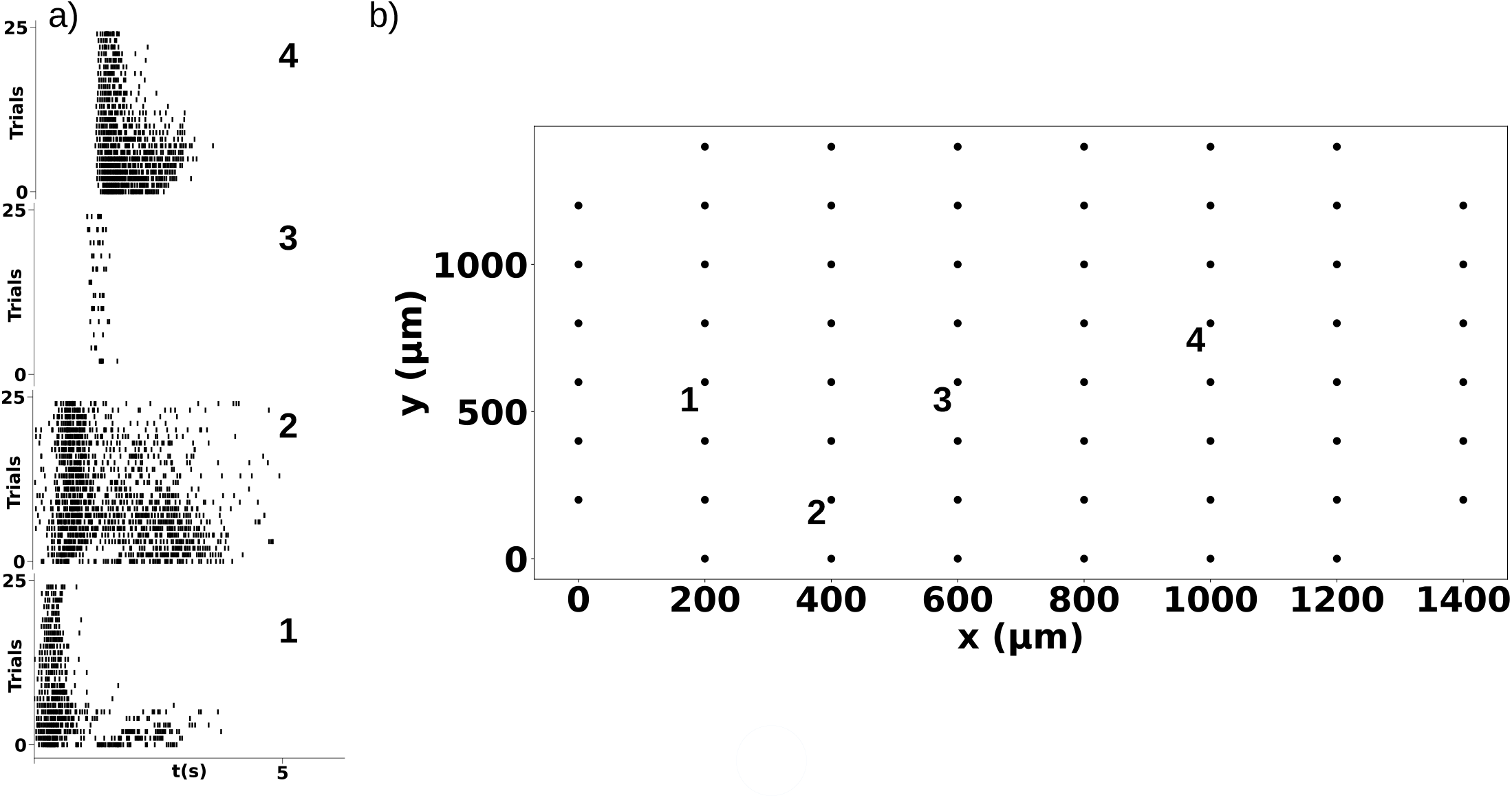
Responses of neonatal RGCs to moving bar stimulus. Response was recorded for bar stimulus moving in the +x direction for a total of 25 trials. a) Raster plots of 4 RGCs showing repeatable responses to the moving bar stimulus. b) Position of the RGCs on the MEA. The time differences in the start of firing shows the direction of bar movement.

The direction of motion of the bar was estimated from differences in the time of peak firing rates of individual RGCs. For each presentation of the moving bar, motion direction was inferred from the relative timing of peak firing across the population of recorded RGCs. Trial-by-trial direction estimates showed consistent clustering around the stimulus direction. The colour maps in Figure 3a)–d) show the time of peak firing of individual RGCs in response to bars moving in different directions across the MEA. As the bar traverses an RGC’s receptive field, its firing rate increases, reaches a maximum, and subsequently decreases. These systematic temporal offsets across the population enabled estimation of motion direction from the collective retinal response (Supplementary Figure 3).

**Figure 3:**
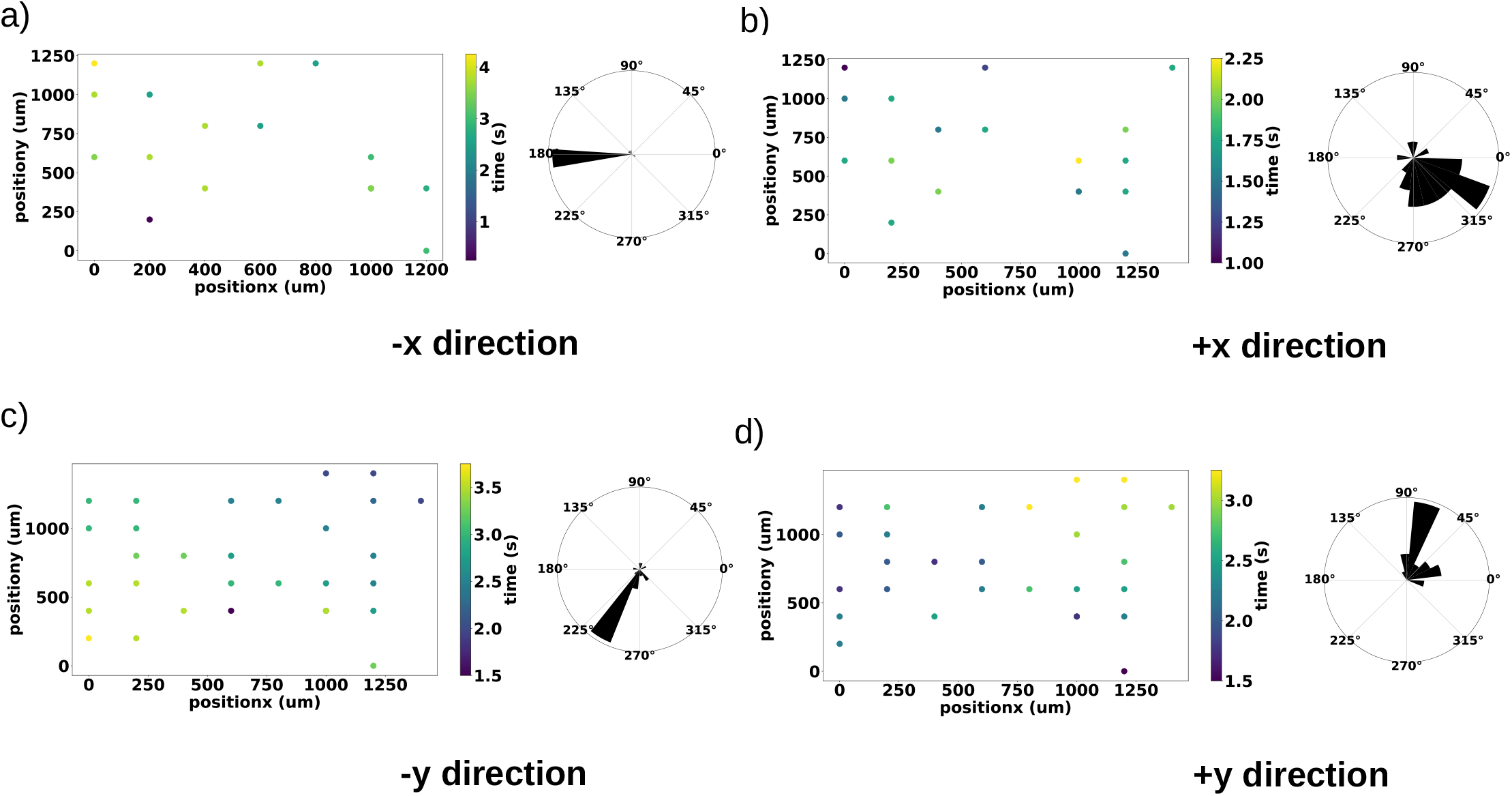
Colour maps depict the trial-averaged time of peak firing rate of identified RGCs across the MEA in response to bar motion in the a) -x, b) +x, c) -y, and d) +y directions. Corresponding rose-plot histograms show the distribution of motion directions estimated on a trial-by-trial basis for bar movement in each direction.

To estimate the speed of the moving bar, we used the temporal differences in peak firing between RGCs situated at different spatial locations. Raster plots of two representative RGCs for varying bar speeds are shown in Figure 4 a). The bar was moved in the -x direction, and the two cells were located at opposite ends of the array (Figure 4 b). The temporal difference in firing onset can be clearly seen in the rasters. As the speed of the bar increased, the rasters came closer, indicating a decrease in relative spike timing. This demonstrates that the speed of the bar can be inferred from the relative spike timing of RGC pairs across spatial locations. Quantitatively, the speed of the moving bar was calculated by using the position of individual RGCs on the MEA and the their time of peak firing. This speed estimate closely matched the actual speed of the bar stimulus (Figure 4 c)), which was determined as the average speed across 20 different pairs of RGCs on the electrode array.

**Figure 4:**
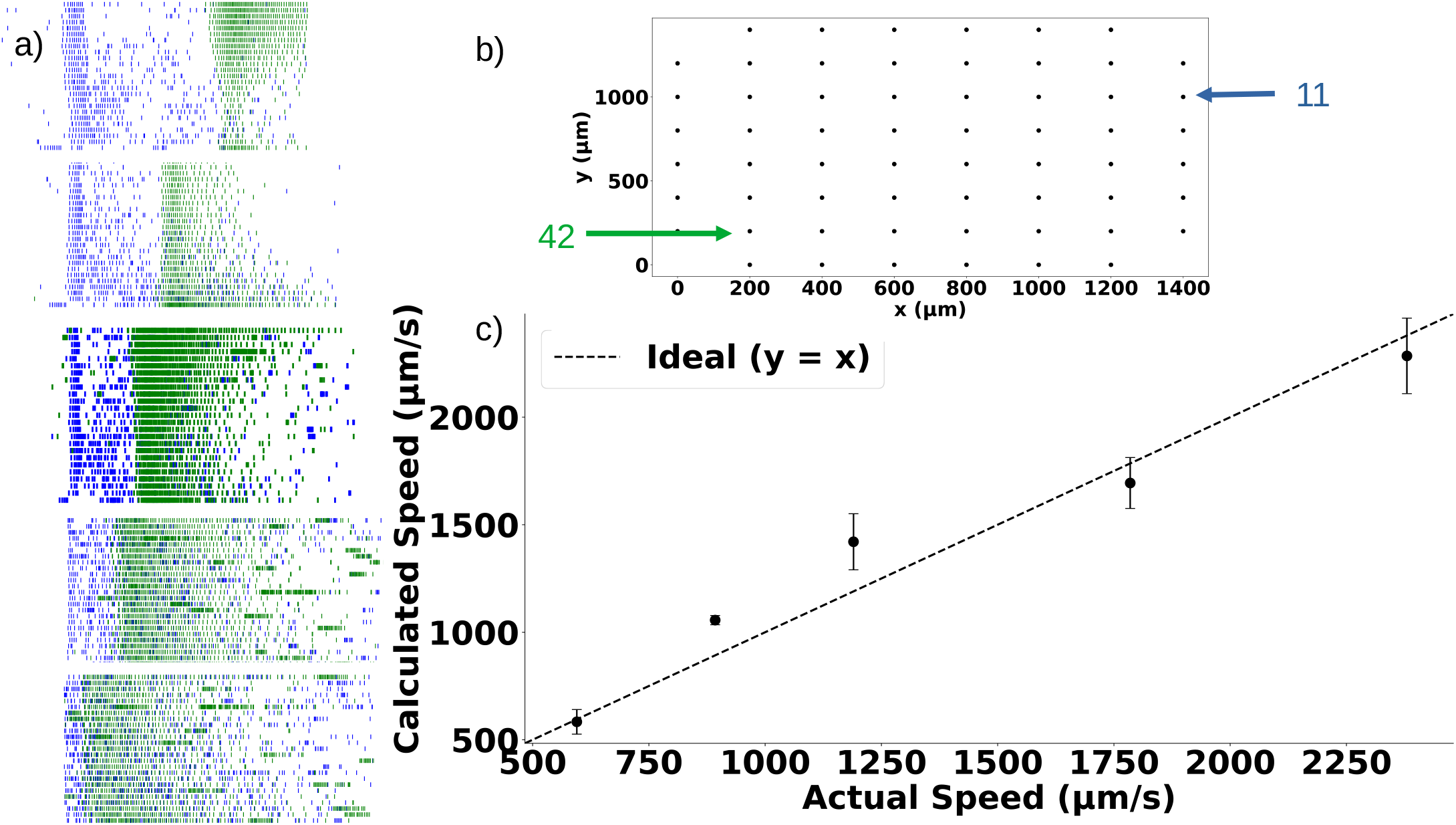
Estimation of the speed of the moving bar using time of peak firing rate and position of RGCs. a) Raster plots from two RGCs (blue and green) for bar movement in the -x direction at different speeds. Speed of the bar increases from top to bottom. b) Position of the two RGCs on the MEA and c)Plot showing the comparison between estimated speed and actual speed in a neonatal retina.

### 2.2 Estimation of direction and speed from prosthetic coupled retina

Subretinal stimulation using photoactive polymer BHJ layer has also been shown to effectively elicit RGC firing[25]. The retina at early stages of development does not show stable light responses due to the lack of a developed photoreceptor layer[14]. Previous studies in the laboratory demonstrated that P3HT:N2200 films can reliably elicit responses from chick RGCs[14, 15]. The prevailing network of bipolars/neurons in the outer retinal layers serves as the conduit for this signal. These unpatterned films that strongly absorbed light reliably activate RGCs both in epiretinal and subretinal configurations, producing ON, OFF and ON–OFF responses.

In the present work, spatially heterogeneous activation was achieved by patterning the P3HT:N2200 films. The solution processible film formation allows soft lithography approaches using Polydimethylsiloxane (PDMS) molds to generate patterns[26]. These patterned films were interfaced subretinally with the developing embryonic chick retina. Pattern widths that are less than few microns were insufficient to reliably elicit RGC activity, likely due to limited charge generation from the small active area and suboptimal contact with the retina[27, 28]. Loss of charges at exposed ITO regions between patterns may also contribute to this unreliable activation[29]. Consequently, the patterned bar width was set at 200 µm, which reliably elicited responses from chick RGCs.

Moving bars of the same width as in the neonatal experiments were projected on to the blind retina interfaced subretinally with the patterned substrate. The patterned films were able to elicit activity from embryonic chick RGCs. Response polarity of RGCs was calculated using full-field flash stimuli with a procedure similar to that as in the neonatal retina. The pulse-width modulation (PWM) of the LEDs in the DLP produced stimulation artifacts when the bar of light passed over the polymer BHJ (Figure 5 a)). These artifacts could be removed during the spike sorting step as they were seen on multiple electrodes. Due to this constraint, only OFF cells identified from full-field flash responses were analysed for direction and speed estimation(Supplementary Figure 4). The moving bar stimulus was repeated over ten trials, and RGCs responded with repeatable firing events, similar to the neonatal case (Figure 5 b) and c)). Evidence of direction-selective RGCs was not clearly observed in the measurements involving polymer-coupled embryonic retina. This can be attributed to the immature amacrine–ganglion cell circuits that are required for such responses during this stage of development.

**Figure 5:**
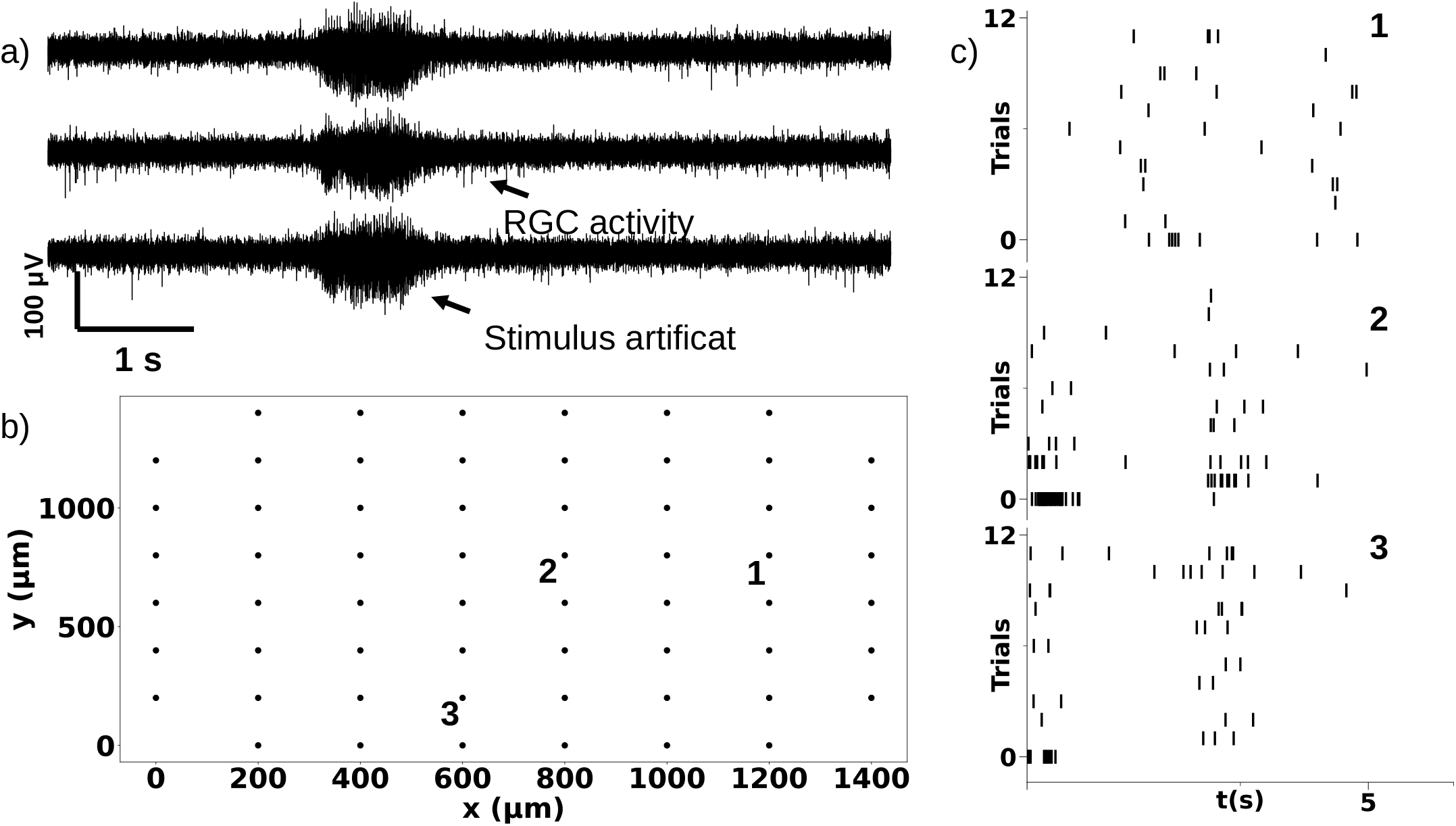
Response of the optically active, semiconducting-polymer coupled retina to moving bar stimuli (-x direction). a) Band-pass filtered data from an embryonic retina subretinally interfaced with the patterned semiconducting polymer layer. The DLP projector used in these experiments produced stimulus artifacts during bar movement. This was removed after spike sorting. b) Positions of the RGCs on the electrode array and c) Raster plots from three RGCs that show temporal delays in their response. b)

The peak firing rate of individual RGCs was calculated from the trial-averaged firing rate. In polymercoupled retinae, temporal differences in peak firing were observed across the RGC population, enabling estimation of the direction of bar motion (Figure 6 a)and b). Due to the width of the patterned P3HT:N2200 bars, activity was elicited on a coarser spatial scale, with RGCs located in adjacent columns often firing simultaneously. Nevertheless, moving bar stimuli projected onto the patterned polymer interface evoked repeatable spatiotemporal firing patterns that could be visualized as colour maps of peak firing time across the electrode array. As the bar traversed the patterned substrate, groups of RGCs exhibited temporally offset peak responses, reflecting sequential activation of RGCs. These population-level temporal differences enabled inference of motion direction in the prosthetic-coupled embryonic retina, indicating that the artificial photoreceptor interface preserves spatiotemporal signatures of object motion. Due to this reduced spatial resolution, the lowest bar speeds used in neonatal retinae were employed in the prosthetic experiments as well, and the estimated speed closely matched the actual stimulus speed (Figure 6 c).

**Figure 6:**
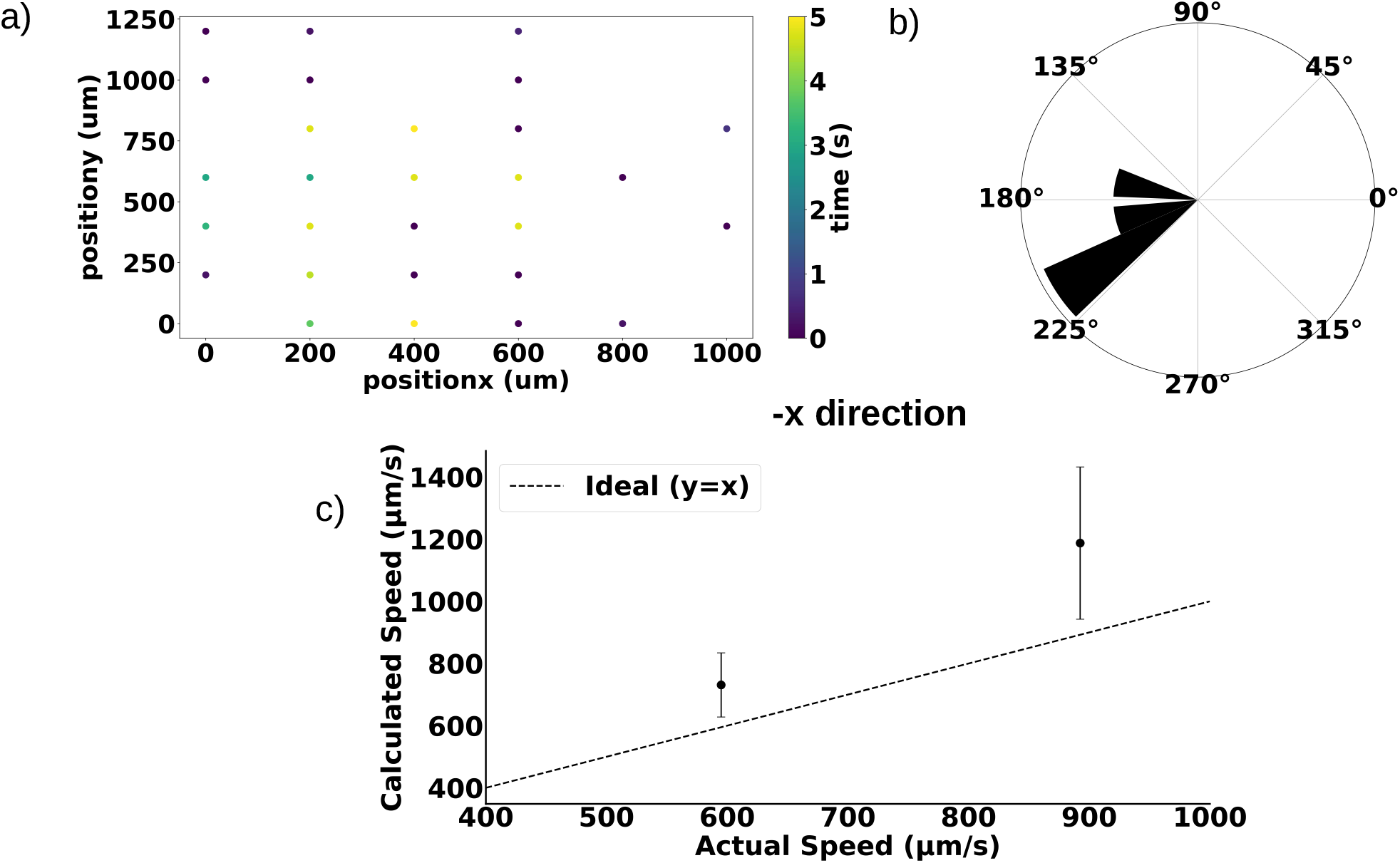
Estimation of the direction and speed of the moving bar using time of peak firing rate and position of RGCs for a prosthetic coupled blind retina. a) Average activity map for bar movement in the -x direction, b) Corresponding rose-plot histogram shows the distribution of motion directions estimated on a trial-by-trial basis for bar movement and c) Calculated speed of the moving bar in a prosthetic coupled retina and its comparison with the actual speed.

A feature of the developing retina is the presence of correlated spontaneous activity known as retinal waves[30]. Our recent studies indicated that the chick retina exhibits propagating spontaneous waves of activity before vision begins[31]. A clear difference in this spontaneous activity was observed in the early embryonic stage compared to the later stages just before hatching. Their presence necessitated limiting the number of trials of the moving bar stimulus to estimate the average peak firing rates, as longer recordings were overwhelmed by a sizable contribution from retinal waves[32]. Despite the presence of retinal waves, the motion parameters could still be reliably extracted from the artificial photoreceptor incorporated retina.

## 3 Conclusion

Initial responses to full-field flashes of light were recorded to establish polarity (ON/OFF cells) of chick RGCs. The response of the RGCs was then studied as a function of the moving bar stimuli. The stimulus dimension was long enough to cover the electrode array along both the axis. The bar was typically swept across the retina along the +x, -x, +y, and -y axes. A subset of RGCs showed direction selective (asymmetric) responses to object motion indicating the presence of DSRGCs in neonatal chick retina. The characteristic firing by a set of DSRGCs can then be used by a downstream layer to identify the absolute direction of the moving bar.

Non-direction selective RGCs responded by firing a burst of spikes when the bar of light moved across its receptive field. This moving stimuli typically results in a visual streak, essentially a wave of activity that spreads across the retina in the direction of bar motion. This travelling wave of activity followed the motion of the bar, and the temporally shifted response from different RGCs can also be used to estimate both the direction and speed of the bar of light. RGCs are known to encode stimulus information through precise spike timings for spatially uniform stimuli, exhibiting low temporal jitter across repeated trials. However, this was not observed in the chick retina for moving spatial stimuli, where RGCs responded with substantial temporal jitter across repeated cycles. This variability makes it unlikely that the timing of the initial spike serves as the primary code for encoding motion direction and speed in the chick retina. Instead, downstream neurons are more likely to rely on firing-rate–based signals from populations of RGCs to estimate object motion. Such a population code offers the advantage that fluctuations in the responses of individual RGCs have minimal impact on the decoded motion parameters, as direction and speed are represented collectively rather than by a single neuron or a small subset of cells, resulting in a motion estimate that is robust to noise. In the present case of the neonatal retina, peak firing rates of RGCs were calculated by averaging responses across individual trials to the moving bar stimulus. Temporal delays between the peak firing rates of RGCs were used to estimate the direction of bar movement. Coupled with the knowledge of RGC positions assigned by spike sorting, these temporal differences also allowed estimation of the speed of the bar.

To understand how downstream neurons might decode these sequential RGC activations, simple computational schemes such as the Reichardt detector are conceptually useful[33]. Such a detector estimates motion by comparing signals from neighbouring spatial locations after introducing a temporal delay to the signal from one spatial location. Motion in one direction causes the delayed and undelayed signals to align, producing a strong output that reflects both the direction and speed of the object. A similar computation may be performed by downstream brain regions in the chick retina which receives spikes from RGCs. The sequential activation of RGCs produced by the moving-bar stimulus naturally provides the temporally offset inputs required for a Reichardt-like detection mechanism.

Earlier work has shown that P3HT:N2200 BHJ layers reliably elicit activity patterns from developing chick RGCs in response to full-field flashes, producing responses similar to those of a healthy (neonatal) retina. In this line of pursuit, studies on the dynamical or motion response were carried out upon introducing the prosthetic layer in the subretinal configuration. This insertion resulted in the activation of outer retinal layers leading to a network mediated activation of RGCs. In the present study, a grating pattern of the P3HT:N2200 film layer, with a 200 µm spacing-width, were interfaced subretinally with the developing chick retina. Motion of a green bar of light elicited a visual streak in the developing blind-retina that resembles the visual streak of the neonatal retina, but with a coarser spatial profile. It was possible to clearly observe the temporal differences in the peak firing rate of RGCs, allowing estimation of motion direction in case of the polymer-coupled retina. It was possible to estimate speed of the bar from the temporally offset responses. This indicated that both direction and speed information is available at a coarse level in a polymer coupled blind retina. However, a clear distinction of specific DSRGCs was not observed over a large set of measurements. This can be attributed to the development stage of the retina and the immaturity of circuits required for such computations.

The technique used in this work to elicit responses from embryonic RGCs can be further optimised. In the present study, the relatively large widths of the patterned polymer bars resulted in activation of RGCs in a larger area (RGCs located on adjacent columns of the MEA) compared to the neonatal recordings. This resulted in the loss of spatial information as RGCs located at different locations were stimulated simultaneously, leading to coarser activity patterns, affecting the precision of speed estimation. Reducing the bar width to improve spatial specificity of activation, resulted in the stimulation of fewer RGCs, making it difficult to determine motion direction. Optimising of both pattern-width and electrode density should enable more accurate estimates of the motion attributes in the prosthetic retina. Nevertheless, it is demonstrated that the important attributes of direction and speed in a vision system can be preserved in an underdeveloped retina using a photoactive polymer layer.

## 4 Experimental Section

### 4.1 Ethics statement

All studies were performed in accordance with the guidelines of the Institutional Animal Ethics Committee (IAEC) and the Institutional Bioethics and Bio-safety Committee, Jawaharlal Nehru Centre for Advanced Scientific Research. All protocols and experiments were approved by the IAEC (Proposal No. KSN 001) and were performed in accordance to the guidelines followed by the Poultry Science Unit, Karnataka Veterinary, Animal and Fisheries Sciences University (KVAFSU, Hebbal, Bangalore).

### 4.2 Device Fabrication

Photoactive polymer bulk heterojunctions (BHJs) were prepared using solution processing. The BHJ blend consisted of P3HT (Luminescence Technology Corp., China) as the electron donor and poly[N,N’-bis(2-octyldodecyl)-naphthalene-1,4,5,8-bis(dicarboximide)-2,6-diyl]-alt-5,5’-(2,2’-bithiophene) (N2200, Polyera Corp., USA) as the electron acceptor. The donor and acceptor polymers were mixed in a 4:1 weight ratio, dissolved in chloro-benzene (Sigma) at a concentration of 10 mg/ml, and stirred overnight to ensure complete dissolution and homogeneous mixing.

The P3HT:N2200 blend was then patterned using soft lithography procedures. A PDMS mold was used to stamp the polymer onto RCA-cleaned indium tin oxide (ITO) coated glass slides of appropriate dimensions. The patterned films were subsequently annealed at 120°C for 15 minutes.

### 4.3 Light Stimuli

Light stimulation was provided using a DLPLCR 4500EVM digital light projector (DLP). The projected light was focused onto the retina through a lens system and a 10× objective of a Nikon inverted microscope, enabling precise control of stimulus location, size and intensity.

For neonatal chick retinae, white light with an intensity range of 100–200 µW/cm^2^ was used to stimulate RGCs. The initial polarity of RGC responses (ON, OFF, or ON–OFF) was determined using 500 ms full-field flashes of light. To probe motion responses, bars of light (200 µm width) were moved across the retina in four directions with respect to the MEA (+x, -x, +y, -y), covering the entire length of the electrode array.

For polymer-coupled embryonic retinae, green light (*λ* = 525 nm) was used to match the peak absorption wavelength of P3HT, with an intensity of 1 mW/cm^2^. The narrow wavelength ensured selective activation of the polymer BHJ while minimizing potential phototoxicity.

### 4.4 Tissue preparation

Retinae were isolated from neonatal (P0–P4) and embryonic (E13–E16) chick eyes following established protocols [15]. Eyes were enucleated under dim light, and the retina was carefully dissected in oxygenated artificial cerebrospinal fluid (aCSF) to preserve tissue viability.

For control recordings, neonatal retinae were placed RGC-side down onto a 60-electrode MEA, allowing direct access of electrodes to the ganglion cell layer for extracellular spike recordings.

For polymer-coupled embryonic retinae, the embryonic retina was similarly placed RGC-side down on the MEA, after which the patterned P3HT:N2200 BHJ film was positioned on top of the photoreceptor layer to create a subretinal interface. This setup allows light-driven activation of the outer retinal network, simulating the function of artificial photoreceptors.

Recordings were performed on four neonatal and four embryonic animals, with each preparation maintained under controlled temperature and oxygenation to ensure tissue health throughout the duration of the recordings.

### 4.5 Direction and speed estimation

For each presentation of the moving bar, the direction of motion was estimated on a trial-by-trial basis from the spatiotemporal pattern of RGC activity. Spike times recorded from individual RGCs were converted to firing rates and smoothened with a Gaussian kernel after which the time of peak firing rate was extracted. The spatial coordinates of the corresponding electrode was used to construct a population activity map. Motion direction was then inferred after fitting with a planar gradient to the time of peak firing across the electrode array, yielding a direction estimate for each trial. Trial-wise direction estimates were visualized using polar histograms (rose plots).

Pairwise spatial separations between RGCs were combined with the temporal differences in their peak firing times to compute local speed estimates. The stimulus speed was then obtained by averaging these pairwise estimates across multiple RGC pairs, yielding a robust population-based measure of motion speed.

## 6 Acknowledgments

This study was supported by the JNCASR-DBT partnership program and JC Bose Fellowship of Department of Science and Technology, India

## 7 Data availability

Data will be made available on request.

