## Supplementory Information for "Estimation of motion direction and speed using an organic-semiconductor retinal prosthetic in a blind retinae"

### Supplementary Information

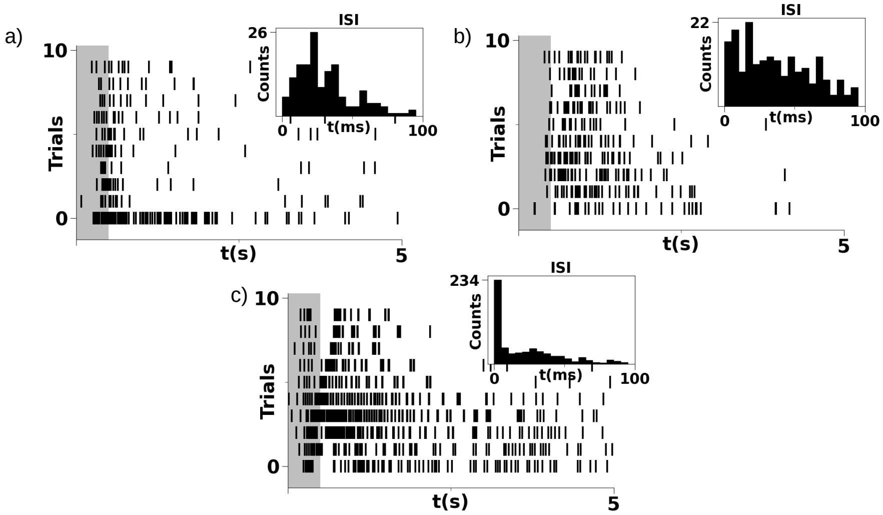


Figure 1: Typical Examples of a) ON, b) OFF and c) ON-OFF RGCs observed in the neonatal chick retina in response to a 500 ms flash of white light.

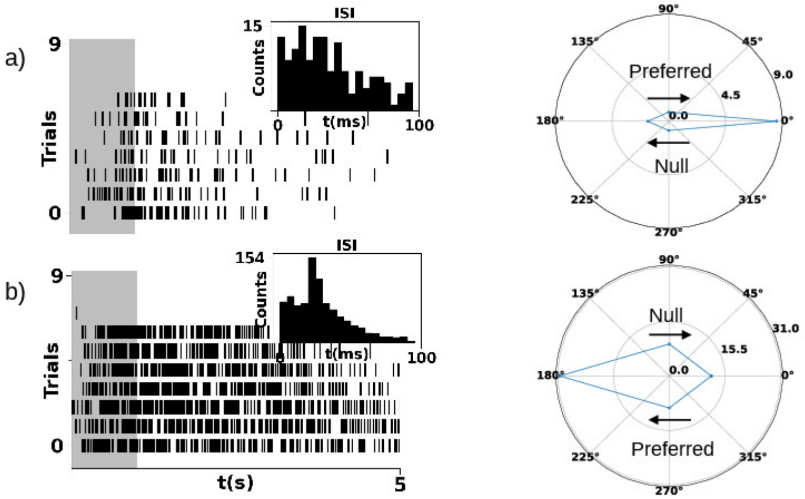


Figure 2: Raster and polar plots of chick RGCs' response to a 500-ms full-field flash and a moving-bar stimulus. a) RGC showing a preferential response to motion in the +x direction and b) RGC showing a preferential response to motion in the -x direction.

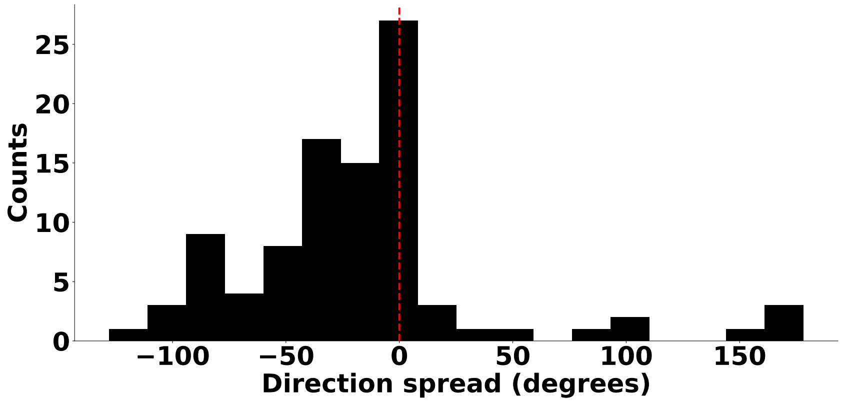


Figure 3: Error range for the estimated direction in the neonatal retina to bar movement. Estimated from results depicted in Figure 3 of the manuscript

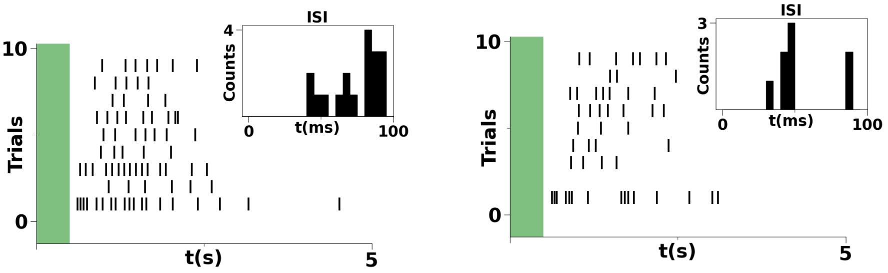


Figure 4: Examples of OFF RGCs seen in the embryonic chick retina subretinally interfaced with the patterned polymer-semiconductor layer in response to a 500 ms flash of green light.
